# Obesity selectively disrupts long-lasting type 2 immune cell depots in white adipose tissue

**DOI:** 10.64898/2026.05.08.723745

**Authors:** Lea Semmler, Andrea Deinzer, Inaya Hayek, Christian Schwartz

**Author notes:** **Correspondence:** Christian Schwartz.

## Abstract

White adipose tissue (WAT) has emerged as a tissue-resident niche for immunological memory, but whether this depot function extends to type 2 immune memory and how it is modulated by metabolic disease remains incompletely understood. Here, we show that *Nippostrongylus brasiliensis* infection, in which the parasite transits and damages the lung, establishes a long-lasting immune cell depot in distal abdominal WAT comprising eosinophils, group 2 innate lymphoid cells (ILC2s), and memory T cells that persists after parasite clearance. Intranasal papain administration recapitulates this depot phenotype, demonstrating that depot formation is a general feature of pulmonary type 2 inflammation rather than a response restricted to live infection. Adoptive transfer of OT-II T cells together with ovalbumin-coated *N. brasiliensis* larvae confirms that WAT accumulates bona fide antigen-experienced T cells. Diet-induced obesity selectively disrupts depot establishment, while leaving acute lung-stage immune cell recruitment, parasite clearance, and the intrinsic competence of cells that do accumulate largely intact. Despite preserved early lung recruitment, obese mice show exacerbated peri-alveolar tissue damage, consistent with an uncoupling of immune cell infiltration from productive type 2-dependent wound repair. Together, these findings establish WAT as a broadly competent type 2 immunological memory niche, identify obesity as a selective disruptor of depot formation rather than effector function, and provide a cellular framework for the impaired vaccine and infection responses associated with obesity.

## Introduction

White adipose tissue (WAT) is an immunologically active organ. Under homeostatic conditions, WAT harbours a characteristic type 2 cell compartment including regulatory T cells (Tregs), eosinophils, alternatively activated macrophages (AAM) and type 2 innate lymphoid cells (ILC2) that maintain each other’s abundance and support metabolic homeostasis [1-3]. Obesity drastically disrupts this balance by also establishing disturbed immune cell functions that may interfere with e.g. T cell polarization [4]. Currently, more than one billion people are living with obesity [5]. Beyond well-established comorbidities such as cardiovascular disease, metabolic dysfunction, and certain types of cancer, obesity profoundly alters immune cell composition, both locally and systemically. In WAT, proinflammatory macrophages [6] and neutrophils [7] accumulate, while eosinophils [8] and ILC2s [9] decline. Comparable shifts have been reported in intestine [10] and liver [11], which collectively lead to increased infection susceptibility and impaired vaccine responses.

Beyond its role in metabolic regulation, WAT has been identified as a reservoir for immunological memory. Han et al. demonstrated that mesenteric adipose tissue accumulates CD69^+^ tissue-resident memory T cells after infection with *Toxoplasma gondii* or *Yersinia pseudotuberculosis*, and that these cells confer protection against secondary infection [12]. Extending this concept to type 2 immunity, Kabat et al. showed that infection with the helminth *Heligmosomoides polygyrus* drives expansion of a type 2-biased resident memory T cells within the mesenteric adipose tissue [13]. As an additional immune cell depot, Collins et al. described the bone marrow as an alternative memory T cell niche during caloric restriction [14].

Notably, these studies examined WAT as a T cell memory depot under lean conditions or dietary restriction. Whether obesity, which fundamentally remodels the WAT immune landscape, impairs depot formation and whether it encompasses long-lived tissue-resident cells other than T cells has not been addressed. We hypothesise that obesity impairs type 2 immune cell depot formation in WAT. To test this, we used the *Nippostrongylus brasiliensis* infection model, which elicits a potent type 2 immune response characterized by eosinophilia, Th2 and ILC2 expansion, and substantial tissue damage within the lung, requiring type 2-dependent wound repair [15, 16]. As type 2 immune responses in general and specifically against *N. brasiliensis* as well as wound healing are affected by obesity [17], we raise the questions, (i) whether *N. brasiliensis* infection establishes a long-term type 2 immune cell depot in WAT, (ii) whether obesity impairs this depot formation during helminth infection and type 2 airway inflammation, and (iii) whether WAT-resident type 2 immune cells are functionally competent to contribute to peripheral immune responses.

## Results

### Helminth infection establishes a long-lasting type 2 immune cell depot in WAT

To confirm and extend previous reports of immune cell depot formation in adipose tissue, we infected wild-type mice maintained on normal chow diet with *N. brasiliensis* and analysed the abdominal WAT at baseline and at 2, 5, 14, and 28 days post infection (p.i.) (Fig. 1A). Unexpectedly, eosinophil and ILC2 numbers declined rapidly within the first days after infection (Fig. 1B, C), while T cell numbers decreased from day 7 until d14 p.i. (Fig. 1D). This early reduction is consistent with redistribution of WAT-resident type 2 immune cells to sites of infection, with the lung being the primary organ of *N. brasiliensis*-induced tissue damage, although direct evidence of trafficking from WAT to lung remains to be established. Strikingly, by d28 p.i., approximately three weeks after parasite clearance, eosinophils, ILC2s, and T cells exceeded their pre-infection abundance in the WAT (Fig. 1B-D), indicating the establishment of a long-lasting type 2 immune cell depot.

**Figure 1.**
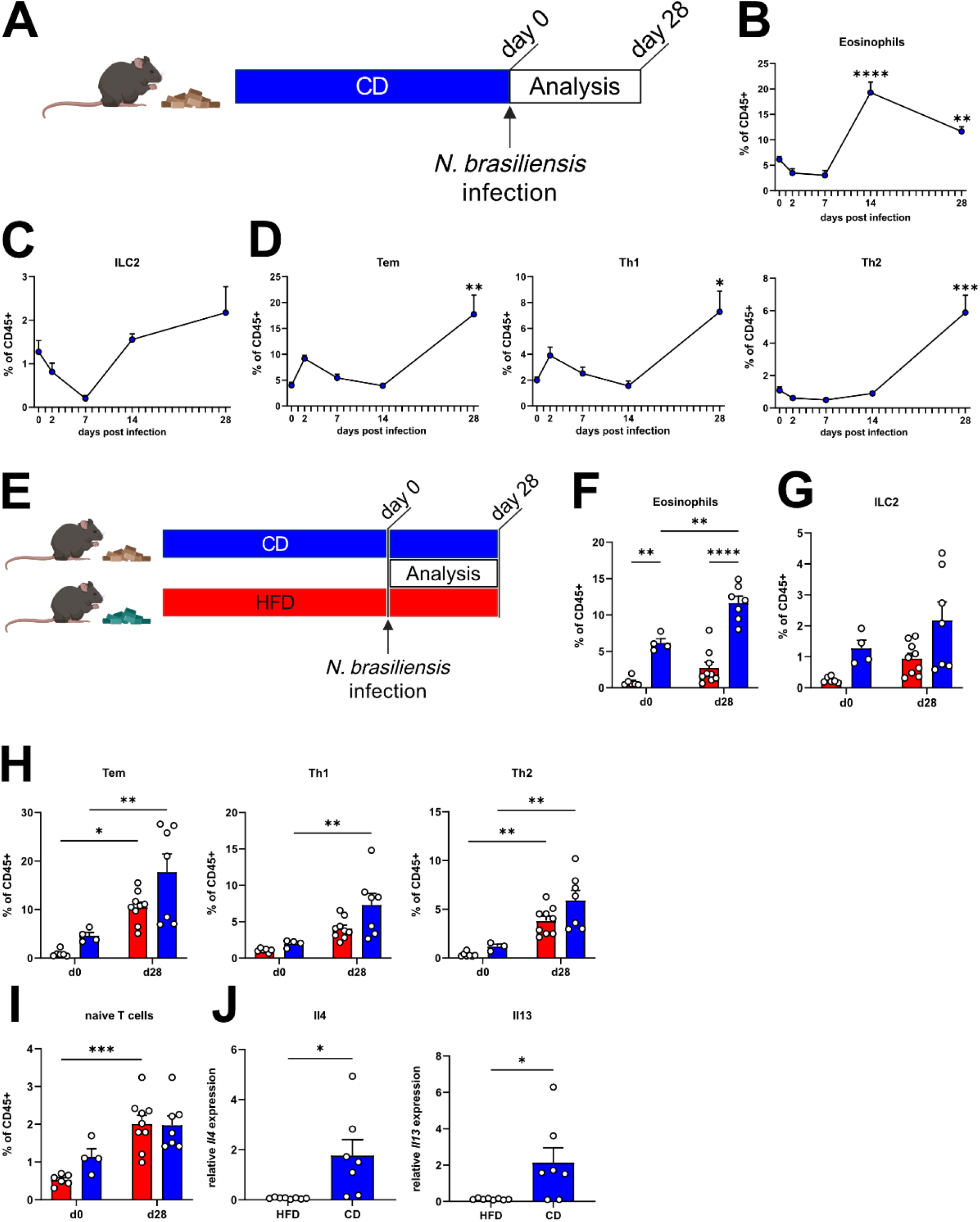
*N. brasiliensis* induces long-lived type 2 immune cell accumulation in lean WAT. **A)** Schematic of mice on control diet (CD) infected with *N. brasiliensis* and analysed for 28 days post infection (d.p.i.). Created with BioRender.com. **B)** Frequency of eosinophils in WAT after *N. brasiliensis* infection. Graph shows mean + SEM of 4-7 mice per group from 3 independent experiments. **C)** Frequency of ILC2 in WAT after *N. brasiliensis* infection. Graph shows mean + SEM of 4-7 mice per group from 3 independent experiments. **D)** Frequencies of effector memory T cells (T_EM_), Th1 and Th2 cells in WAT after *N. brasiliensis* infection. Graphs show mean + SEM of 4-7 mice per group from 3 independent experiments. **E)** Schematic of mice on high fat diet (HFD, red) and control diet (CD, blue) infected with *N. brasiliensis* and analysed at 28 d.p.i.. **F)** Frequency of eosinophils in WAT of naïve animals (d0) and at d28 post *N. brasiliensis* infection. Bars show mean + SEM of 4-9 animals per group from 3 independent experiments. **G)** Frequency of ILC2 in WAT of naïve animals (d0) and at d28 post *N. brasiliensis* infection. Bars show mean + SEM of 4-9 animals per group from 3 independent experiments. **H)** Frequency of effector memory T cells (T_EM_), Th1 and Th2 cells in WAT of naïve animals (d0) and at d28 post *N. brasiliensis* infection. Bars show mean + SEM of 4-9 animals per group from 3 independent experiments. **I)** Frequency of naïve T cells in WAT of naïve animals (d0) and at d28 post *N. brasiliensis* infection. Bars show mean + SEM of 4-9 animals per group from 3 independent experiments. **J)** Relative expression of *Il4* and *Il13* (normalized to *Hprt1*) in WAT of *N. brasiliensis*-infected animals at d28 p.i. comparing obese and lean WAT. Bars show mean + SEM of 7-8 animals per group from two independent experiments. \**p* < 0.05, \*\**p* < 0.01, \*\*\*\**p* < 0.0001; one-way ANOVA (B-D), two-way ANOVA (F-I) or unpaired t-test.

### Obesity impairs type 2 immune cell depot formation in WAT

To determine whether obesity affects depot formation, mice were maintained on high-fat diet (HFD) or control diet (CD) for at least 12 weeks prior to *N. brasiliensis* infection. WAT was analysed at baseline and d28 p.i. (Fig. 1E-J, Supplemental Fig. 1). While CD-fed mice showed robust eosinophil accumulation following infection, this response was diminished in HFD-fed mice (Fig. 1F). ILC2 expansion showed a non-significant trend toward attenuation under HFD compared to CD conditions (Fig. 1G). Importantly, T cell accumulation was markedly more pronounced in CD-fed than in HFD-fed mice (Fig. 1H), suggesting that obesity impairs the long-term maintenance of the type 2 immune cell depot in WAT. Accumulation of naïve T cells on day 28 after infection was not altered between mice on CD and HFD, suggesting selective memory disruption by obesity (Fig. 1I). Consistent with the defect in cell accumulation, qPCR analysis of WAT at d28 p.i. revealed reduced expression of the type 2 effector cytokines *Il4* and *Il13* in HFD-fed mice (Fig. 1J).

### Obesity does not impair early lung immune cell recruitment but exacerbates pulmonary tissue damage

To assess whether obesity alters the acute immune response at the primary site of infection, we analysed lung immune cell composition at d7 p.i. in CD- and HFD-fed mice. Total lung cellularity was comparable between groups (Fig. 2A). Eosinophils showed pronounced infiltration into the lungs by d7 p.i., with similar accumulation in both dietary conditions (Fig. 2B). Neutrophil frequencies declined by d7, consistent with their role in the early innate response preceding eosinophil recruitment (Fig. 2B). AAM and ILC2 frequencies were likewise comparable between CD- and HFD-fed mice (Fig. 2C, D). To confirm that parasites successfully reached the lungs regardless of body composition, we quantified larvae at d2 p.i. and found equivalent worm burdens in the lungs of both groups (Fig. 2E), ruling out impaired larval migration. Despite comparable immune cell recruitment, histological analysis of HFD lungs revealed increased granularity and signs of haemorrhage at d7 p.i., while CD lungs showed largely restored architecture (Fig. 2F), suggesting that obesity may delay tissue repair independently of immune cell infiltration.

**Figure 2.**
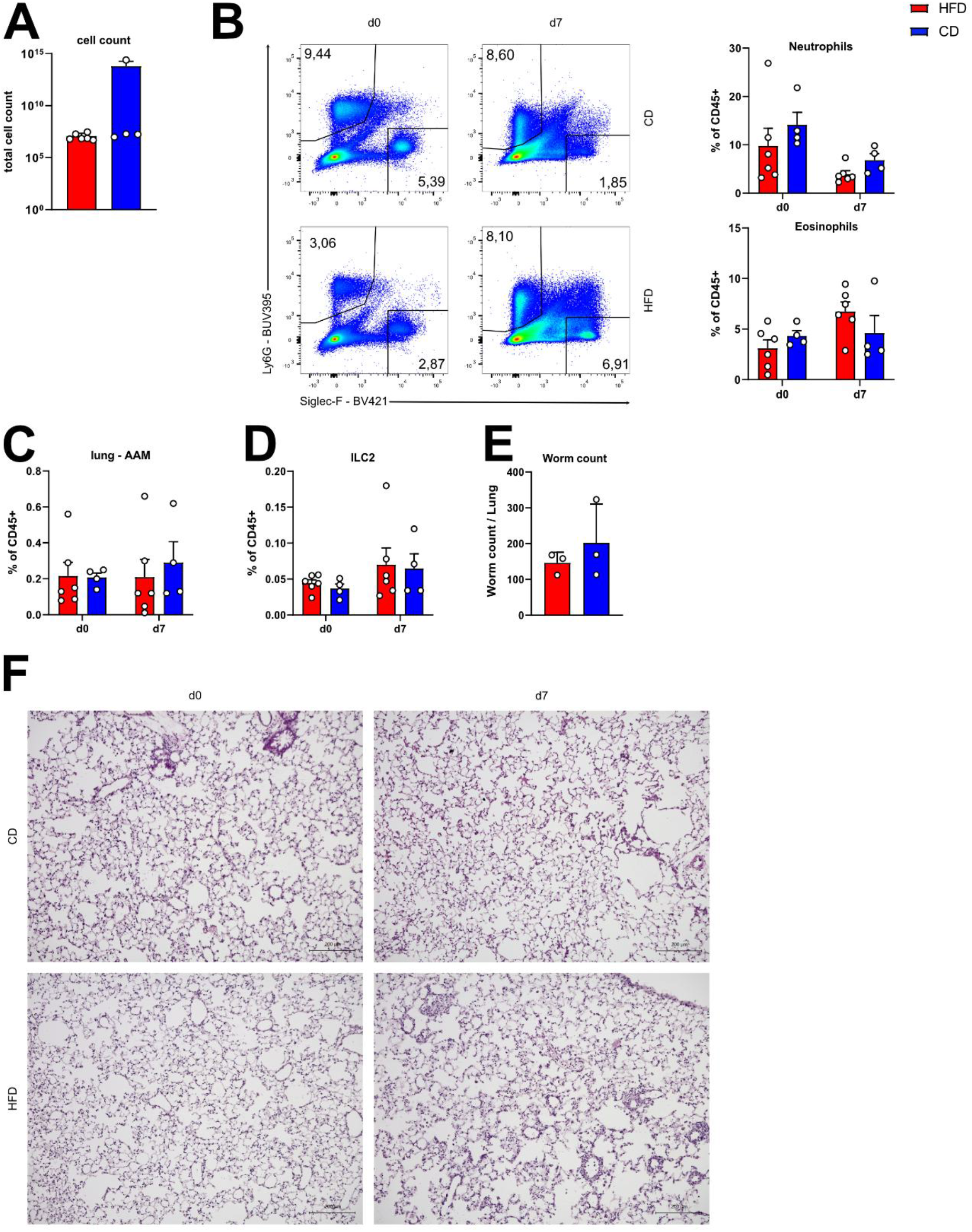
Type 2 immune response to *N. brasiliensis* infection in the lung is largely preserved. **A)** Total cell counts in the lung of obese (red) and lean (blue) animals at d7 post *N. brasiliensis* infection. Bars show mean + SEM of 4-6 animals. **B)** Representative flow cytometric plots and quantification of eosinophils and neutrophils in lung samples of lean and obese animals before (d0) and 7 days post *N. brasiliensis* infection (d7). Bars show mean + SEM of 4-6 animals per group from 2 independent experiments. **C)** Frequency of alternatively activated macrophages (AAM) in the lung of naïve animals (d0) and d7 post *N. brasiliensis* infection. Bars show mean + SEM of 4-6 animals per group from 2 independent experiments. **D)** Frequency of ILC2s in the lung of naïve animals (d0) and d7 post *N. brasiliensis* infection. Bars show mean + SEM of 4-6 animals per group from 2 independent experiments. **E)** Total worm count in the lung of obese and lean animals 2 days post *N. brasiliensis* infection. Bars show mean + SEM from 3 animals per group. **F)** Representative histological sections of naïve (d0) and *N. brasiliensis*-infected (d7) lungs stained with standard haematoxylin and eosin staining. Magnification: 10x; scale bars: 200 µm.

### Allergic airway inflammation recapitulates obesity-dependent impairment of WAT depot formation

To determine whether the observed WAT depot formation extends beyond infectious stimuli, we employed intranasal papain administration as a model of allergic airway inflammation. Mice maintained on CD or HFD were sensitised on three consecutive days and analysed at d5, 7, 10, 14, and 21 after the last treatment (Fig. 3A). As expected, papain induced robust airway inflammation in both groups, evidenced by marked cellular infiltration into the lung parenchyma peaking at d7. More pronounced and diffuse bleeding was observable in HFD-fed animals compared to CD-fed animals (Fig. 3B).

**Figure 3.**
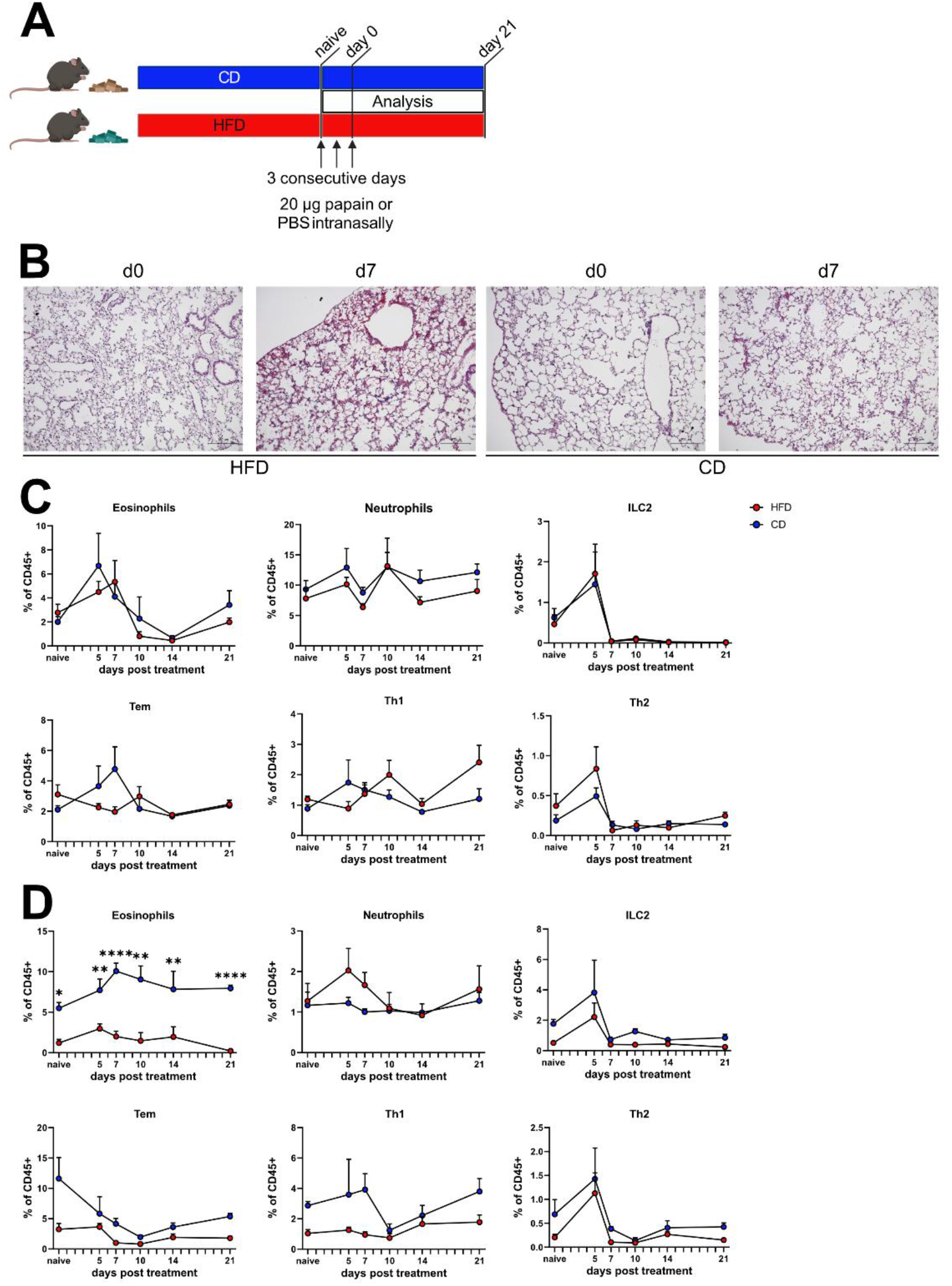
WAT depot formation is independent of infection. **A)** Schematic of mice on high fat diet (HFD) and control diet (CD) treated with 20 µg papain intranasally on three consecutive days and analysed afterwards for 21 days. **B)** Representative histological sections of naive (d0) and papain-treated (d7) lungs from lean and obese animals. Sections were stained with standard haematoxylin and eosin staining. Magnification: 10x; scale bars: 200 µm. **C)** Frequencies of eosinophils, neutrophils, ILC2s, effector memory T cells (T_EM_), Th1 and Th2 cells in the lung of lean (blue) and obese (red) mice at indicated days after papain treatment. Graphs show mean + SEM of up to 6 mice per group of 2 independent experiments. **D)** Frequencies of eosinophils, neutrophils, ILC2s, effector memory T cells (T_EM_), Th1 and Th2 cells in the WAT of lean (blue) and obese (red) mice at indicated days after papain treatment. Graphs show mean + SEM of up to 6 mice per group of 2 independent experiments. \**p* < 0.05, \*\**p* < 0.01, \*\*\*\**p* < 0.0001; two-way ANOVA.

Detailed immune cell kinetics in the lung revealed largely comparable responses between CD- and HFD-fed mice, with some notable exceptions. Eosinophils accumulated in the lungs of both groups, with HFD-fed mice experiencing a minor delay in recruitment. In both conditions, eosinophil numbers returned to baseline by d14 (Fig. 3C). Neutrophil frequencies remained relatively stable throughout, consistent with the neutrophil-independent nature of this type 2 airway inflammation model (Fig. 3C). ILC2 numbers peaked at d5 and subsequently declined sharply to below baseline levels, irrespective of diet (Fig. 3C). Among T cell subsets, effector memory T cells increased during the acute phase exclusively in CD-fed mice. Th1 cells fluctuated around baseline, while Th2 cells followed kinetics similar to ILC2s - peaking at d5 before sharply declining - independent of dietary condition (Fig. 3C). Mirroring our findings after *N. brasiliensis* infection, analysis of abdominal WAT revealed diet-dependent differences in type 2 immune cell accumulation. Eosinophils, which were already reduced at baseline in HFD-fed mice consistent with obesity-driven depletion (Fig. 1F), increased further in the WAT of CD-fed mice following papain treatment and, critically, remained elevated above pre-treatment levels three weeks after cessation of airway challenge (Fig. 3D). In contrast, HFD-fed mice showed no eosinophil accumulation in WAT at any time point examined, neither during the acute phase nor at d21 (Fig. 3D). Of note, neutrophils transiently increased in the WAT of HFD-fed mice only (d5-7), potentially reflecting the pro-inflammatory milieu of obese adipose tissue (Fig. 3D).

ILC2s increased transiently in the WAT of both CD- and HFD-fed mice following papain treatment, peaking at d5 before returning to baseline, with no evidence of sustained elevation in either group (Fig. 3D). T cell dynamics in the WAT were more complex. Effector memory T cells were more abundant in naive CD-fed than HFD-fed mice at baseline. Following papain treatment, they declined in both groups before partially recovering by d21 in CD-fed but not HFD-fed mice (Fig. 3D). Th1 cells followed similar kinetics, while Th2 cells accumulated in both groups until d5 before returning to baseline (Fig. 3D). These results suggest that acute responses remain intact, while long-term depot formation is impaired by obesity.

Together, these data demonstrate that type 2 immune challenge of the airways even in the absence of infection, triggers WAT immune cell dynamics that mirror those observed after helminth infection. Eosinophils appeared to be consistently accumulating in the WAT of CD-fed mice, maintaining elevated numbers well beyond resolution of airway inflammation. T cell responses in the WAT were more transient in the papain model, though their contribution to depot formation and maintenance cannot be excluded. Importantly, obesity impaired WAT depot formation across both inflammatory contexts, reinforcing a general vulnerability of type 2 immune memory in adipose tissue under metabolic stress.

### Antigen-specific T cells accumulate in the WAT depot following helminth infection

The accumulation of T cells in WAT following *N. brasiliensis* infection raised the question whether these cells are antigen-specific or bystander populations recruited non-specifically during inflammation. To address this, we adoptively transferred 6 x 10^6^ OT-II transgenic CD4^+^ T cells, which express a TCR specific for OVA_323-339_, into naive recipient mice. One day after transfer, mice were infected with *N. brasiliensis* pre-incubated with ovalbumin (Fig. 4A). At day 7 p.i., a distinct OT-II population was detectable in both lung and WAT of recipient animals (Fig. 4B). OT-II cells were not recovered from either tissue in the absence of infection, confirming that antigen-specific activation during infection is required for their expansion and tissue accumulation. When comparing dietary conditions, CD-fed mice showed a trend toward higher frequencies of antigen-specific T cells in the WAT following infection, though variability was high and sample sizes limited, precluding definitive conclusions at this stage (Fig. 4C). These findings demonstrate that the WAT depot contains bona fide antigen-experienced T cells, rather than merely reflecting passive redistribution of lymphocytes during systemic inflammation.

**Figure 4.**
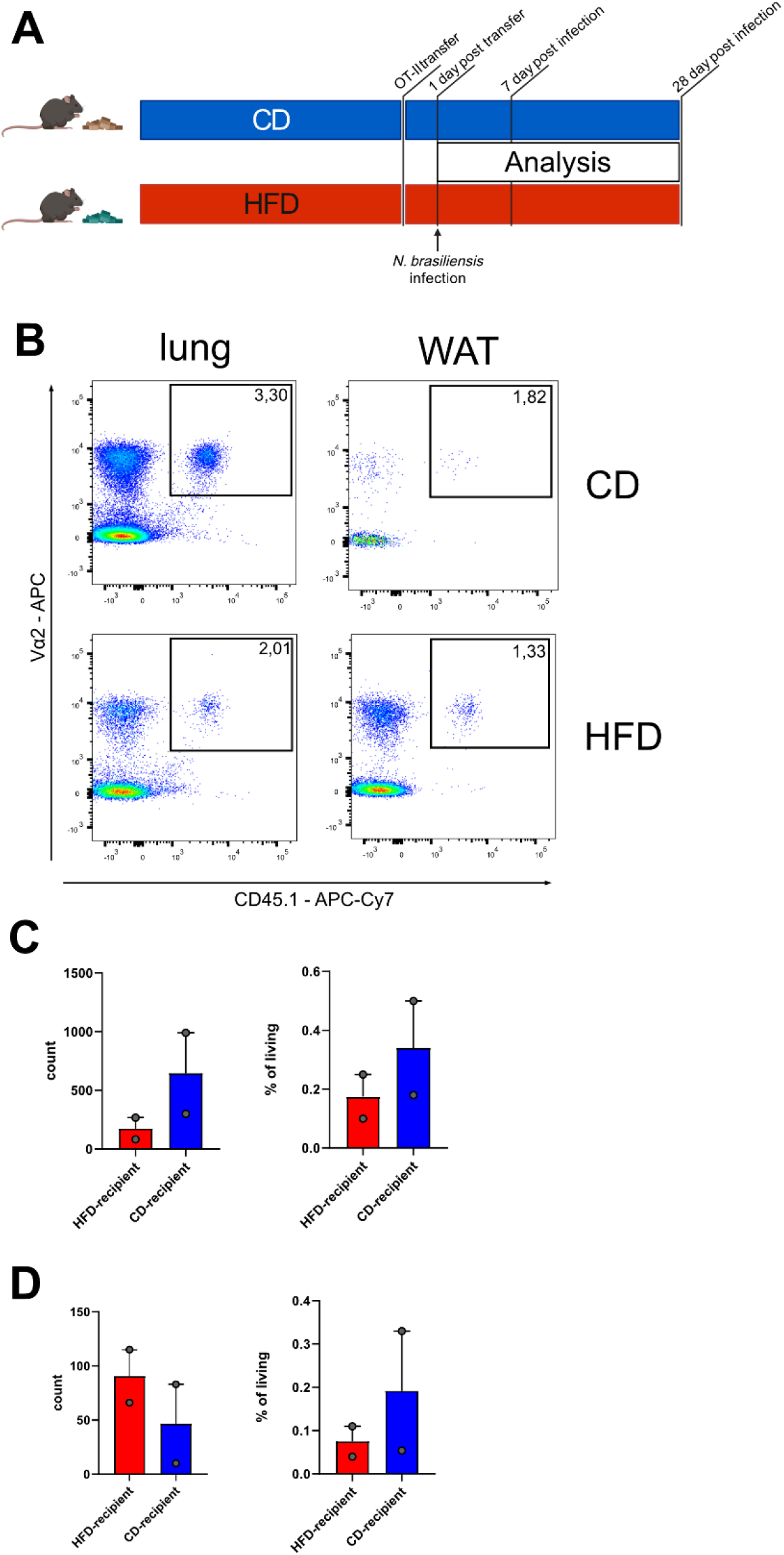
WAT accumulates antigen-specific cells, which partake in the immune response. **A)** Schematic of the OT-II CD4^+^ T cell transfer. Lean (blue) or obese (red) mice received 6 x 10^6^ OT-II CD4^+^ T cells from the spleen of *Tg(TcraTcrb)425Cbn/J CD45*.*1* mice (donors). Recipient animals were infected with *N. brasiliensis* in combination with ovalbumin (OVA) 1 day post transfer and analysed at 7 days post infection (d.p.i.). **B)** Representative FACS plots from lung and WAT from lean and obese animals at 7 days post infection displaying CD45.1^+^ and Vα2^+^ transferred immune cells. **C)** Quantification and frequency of transferred OT-II cells among live cells in the lung of CD-fed and HFD-fed recipient animals at 7 d.p.i.. Bars show mean + SEM of 2 animals per group. **D)** Quantification and frequency of transferred OT-II cells in the WAT of CD-fed and HFD-fed recipient animals at 7 d.p.i.. Bars show mean + SEM of 2 animals per group.

### Transfer of WAT-resident immune cells contributes to pulmonary anti-helminth responses

We next asked whether the type 2 depot immune cells are functionally competent and can participate in a secondary immune response. To this end, we isolated total immune cells from the WAT of Ly5.1 congenic mice at d28 after *N. brasiliensis* infection from mice kept on CD or HFD, and transferred them into *Rag2*^*-/-*^*Il2rg*^*-/-*^ recipient mice, which lack endogenous T, B, and innate lymphoid cells. Recipients were infected with *N. brasiliensis* one day after transfer and analysed at d7 p.i. (Fig. 5A). Donor-derived T cells, including effector memory, Th1, and Th2 subsets, were recovered from the lungs of recipient animals in both CD and HFD transfer groups (Fig. 5B, C). Notably, T cell expansion occurred predominantly in infected recipients that received *N. brasiliensis*-experienced WAT-derived cells, indicating antigen-driven re-activation of the transferred depot cells rather than homeostatic proliferation alone.

**Figure 5.**
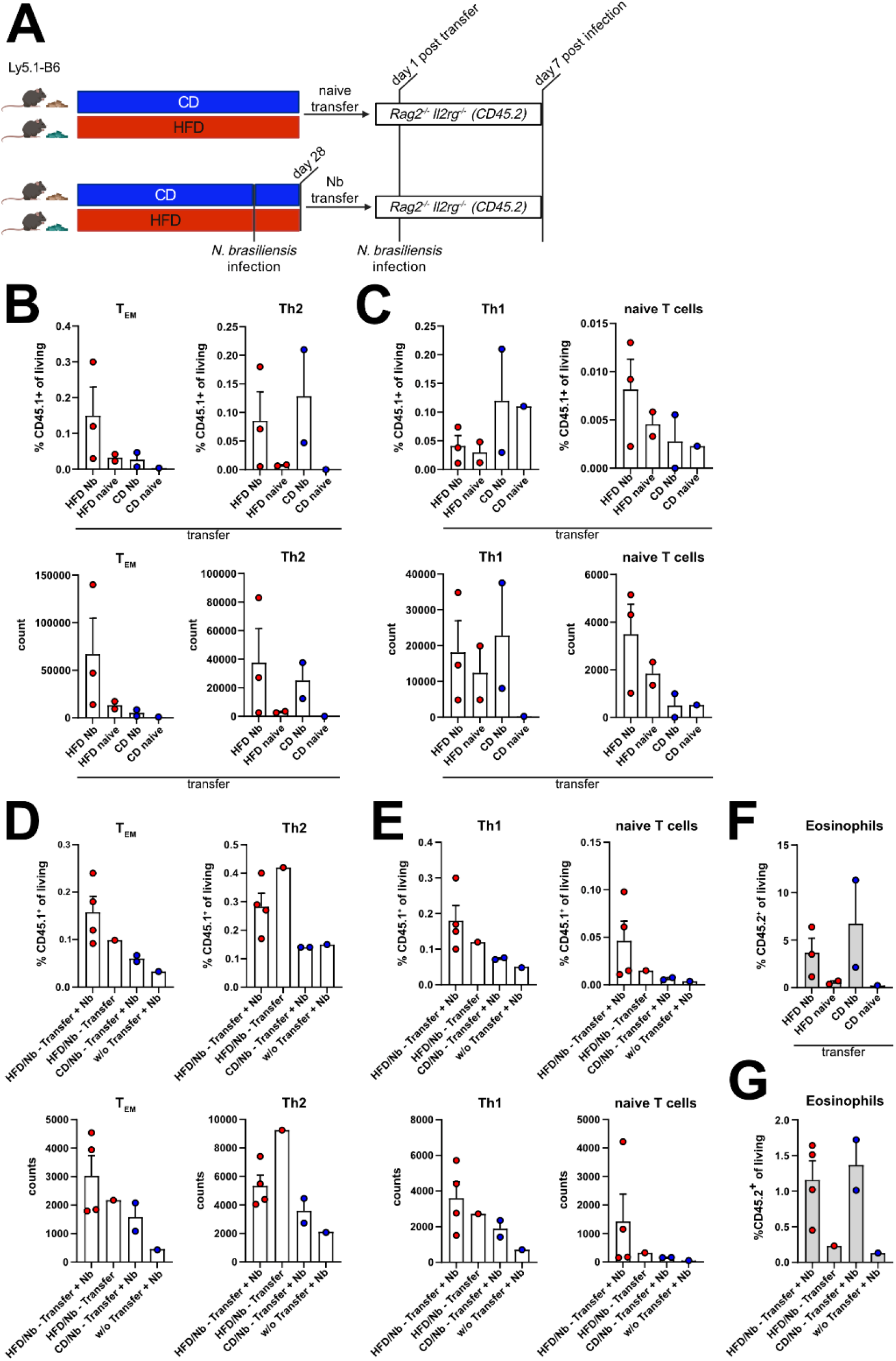
Pre-experienced transferred WAT immune cells migrate to site of infection. **A)** Schematic of the WAT immune cell transfer. Lean (blue) and obese (red) Ly5.1-animals (donors) were either infected with *N. brasiliensis* for infected transfers or kept naïve for naïve transfers. Either total immune cells from the WAT or CD4^+^ T cells from the WAT were transferred into *Rag2*^*-/-*^ *Il2rg*^*-/-*^ (CD45.2) mice (recipients), which were infected with *N. brasiliensis* one day post transfer and analysed at 7 days post infection (d.p.i.). **B)** Frequencies and quantification of transferred effector memory T cells (T_EM_) and Th2 cells in the lung of recipient animals at 7 d.p.i. after total WAT immune cell transfer. Bars show mean + SEM of 2 independent experiments. **C)** Frequencies and quantification of transferred Th1 and naïve T cells in the lung of recipient animals at 7 d.p.i. after total WAT immune cell transfer. Bars show mean + SEM of 2 independent experiments. **D)** Frequencies and quantification of transferred effector memory T cells (T_EM_) and Th2 cells in the lung of recipient animals at 7 d.p.i. after CD4^+^ T cell transfer. Bars show mean + SEM. **E)** Frequencies and quantification of transferred Th1 and naïve T cells in the lung of recipient animals at 7 d.p.i. after CD4^+^ T cell transfer. Bars show mean + SEM. **F)** Frequency of recipient intrinsic eosinophils at 7 d.p.i. after total WAT immune cell transfer. Bars show mean + SEM from 2 independent experiments. **G)** Frequency of recipient intrinsic eosinophils at 7d.p.i. after CD4^+^ T cell transfer. Bars show mean + SEM.

To more specifically assess the T cell contribution, we magnetically isolated CD4^+^ T cells from WAT depots and transferred them into *Rag2*^*-/-*^*Il2rg*^*-/-*^ recipients. Purified T cells preferentially accumulated in the lungs following *N. brasiliensis* infection compared to uninfected recipients, confirming active recruitment to the site of infection (Fig. 5D, E). However, no difference was observed between cells derived from CD-fed and HFD-fed donors after infection, suggesting that once T cells have been established in the WAT depot, their capacity to respond to secondary challenge may be preserved regardless of the metabolic environment of origin (Fig. 5D, E). Importantly, transfer of either total WAT cells or purified CD4^+^ T cells was sufficient to elicit expansion of endogenous eosinophils in the lungs of recipient mice following infection (Fig. 5F, G).

Collectively, these data establish that *N. brasiliensis* infection generates a long-lasting, antigen-specific type 2 immune cell depot in WAT that is functionally capable of contributing to secondary immune responses. Obesity selectively impairs the formation of this depot but does not appear to compromise the intrinsic functionality of the immune cells that do accumulate.

## Discussion

Adipose tissue has emerged as a tissue-resident niche for immunological memory, with prior work establishing roles in CD8^+^ memory responses to intracellular pathogens [12] and in mesenteric WAT after enteric helminth infection [13], paralleling the lung Th2-polarized tissue-resident memory cell findings [18]. Here, we extend this concept in three directions. First, we demonstrate that *N. brasiliensis* infection, in which the parasite predominantly transits and damages the lung, establishes a long-lasting immune cell depot in distal abdominal WAT, indicating that depot formation is not restricted to anatomical proximity between the niche and the primary site of infection. Second, we show that this depot harbours not only memory T cells but also tissue-resident innate cells [19], including eosinophils and ILC2s, which persist for at least three weeks after parasite clearance. Third, we identify obesity as a selective disruptor of depot establishment, while leaving acute lung-stage immune cell recruitment and the intrinsic functional competence of cells that do accumulate largely intact. These findings extend our prior demonstration that innate PD-L1 limits T cell-mediated WAT inflammation under HFD [4] and offer a cellular framework for impaired vaccine responses and accelerated antibody waning observed in obese individuals across pathogens [20].

Eosinophils are generally regarded as short-lived effector cells, which show prolonged survival following helminth infection [21], mediated by IL-5, GM-CSF, and IL-3 [22]. Adipose tissue eosinophils represent a noteworthy exception, being maintained at homeostasis by ILC2-derived IL-5 and contributing to AAM abundance and metabolic regulation [1, 8]. Our findings indicate that infection-driven type 2 inflammation either expands this resident eosinophil pool or seeds a long-lived population that exceeds homeostatic numbers and persists weeks beyond parasite clearance. Adoptive transfer of young adipose eosinophils into aged hosts rejuvenates WAT immune fitness [23], underscoring functional heterogeneity. A recent cross-tissue atlas reveals striking compartment-specific differences in eosinophil longevity: lung eosinophils are short-lived while intestinal eosinophils persist for weeks [24]. Where infection-expanded gonadal WAT eosinophils sit on this continuum remains an open question; whether infection-driven type 2 inflammation expands a long-lived resident pool, seeds a phenotypically distinct subset, or relies on continuous bone-marrow replenishment will require fate-mapping approaches that single-cell profiling alone cannot resolve.

Importantly, depot formation seemed to be independent of the pulmonary challenge. Papain administration led to similar WAT depot kinetics compared to live helminth infection, suggesting a general feature of type 2 inflammation. Critically, the antigen-specific T cell and innate ILC2 components of the depot likely reflect mechanistically distinct memory programmes. IL-25 alone, without cognate antigen, has recently been shown to induce long-lasting memory ILC2s via epigenetic reprogramming [25], establishing cytokine-driven innate memory as a paradigm parallel to adaptive Th2 resident memory in the lung [18]. The papain depot, which forms independently of cognate antigen, fits this innate cytokine-imprinted paradigm, while OT-II accumulation following co-administration of *N. brasiliensis* and ovalbumin establishes that antigen-specific Th2 seeding occurs in parallel during live infection. Whether ILC2-OX40L bridges these two arms by licensing adaptive Th2 expansion within the depot, as shown for primary helminth responses [26], remains to be determined.

Several mechanisms may underlie the obesity-driven depot defect. Obese WAT exhibits a proinflammatory cytokine landscape, including IFNγ accumulation that constrains type 2 lymphocyte niches under mixed inflammation [17] and counter-regulates WAT ILC2 maintenance via IL-33-IFNγ antagonism [27]. HFD-induced loss of LKB1 has recently been identified as a driver of PD-1^+^ exhausted ILC2s in adipose tissue [28], with PD-1 directly destabilizing obese WAT ILC2s [29] which reflect our previous data that innate PD-L1 limits T cell-mediated WAT inflammation under HFD [4]. Diet-induced expansion of pro-inflammatory macrophages and neutrophils [6, 7] alongside baseline loss of resident eosinophils [8] and ILC2s [9] further erodes the type 2-supportive milieu required for depot stabilization. Consistent with a functional correlate, qPCR analysis of WAT at d28 p.i. (Fig. 1J) revealed reduced expression of type 2 effector and tissue-remodelling genes in HFD-fed mice.

Strikingly, WAT-derived immune cells from HFD-fed donors transferred into lymphocyte-deficient recipients trafficked to the lung and contributed to anti-helminth responses comparably to CD-derived cells, paralleling reports that adoptively transferred eosinophils can functionally rejuvenate aged tissues [23]. This dissociation between depot establishment and intrinsic cell competence suggests that obesity primarily acts at the level of niche formation and recruitment rather than corrupting memory function once established. However, a study by Misumi et al. shows that obesity expands a phenotypically distinct antigen-specific WAT T cell population which, upon viral rechallenge, drives immunopathology rather than protection [30]. The discrepancy likely reflects fundamental differences between type 2 and type 1 memory: type 2 effector programmes may be less susceptible to obesity-induced functional rewiring, or the helminth recall readout may not capture more subtle dysfunctions apparent only against chronic viral antigen. The dissociation we observe nonetheless implies that obesity-associated immunological vulnerability may be most pronounced at priming and depot formation, raising the possibility that preinfection weight loss intervention could partially rescue impaired memory responses, as previously shown for vaccine-induced immunity [31].

Although total immune cell recruitment to the lung at d7 p.i. was comparable between dietary conditions, pulmonary tissue damage was exacerbated in HFD-fed mice. Equivalent worm burden in the lungs at d2 p.i. exclude differences in larval migration, and comparable eosinophil, ILC2, and AAM accumulation argue against a recruitment defect. Increased peri-alveolar cell infiltration in HFD lungs is therefore most consistent with delayed resolution of inflammation and wound healing, in line with the dependence of *N. brasiliensis*-induced lung repair on type 2 cytokine-driven processes [15] and on AAM-derived RELMα and Ym1 that limit IL-17-mediated immunopathology and emphysema [16, 32]. Notably, ILC2-derived AMCase has recently been shown to drive epithelial repair after lung injury, with its loss producing a fibrotic phenotype reminiscent of that observed here in HFD lungs [33]. HFD has independently been shown to impair alveolar epithelial repair after bleomycin [34] and influenza-induced lung wound healing despite comparable viral load [35], supporting the interpretation that obesity uncouples immune cell recruitment from productive tissue repair.

Several limitations of our study should be considered. Our analyses focus on gonadal WAT in male mice, leaving open whether sex-dimorphic WAT immune imprinting, which includes a female-enriched CXCR3^+^ Treg population that inversely correlates with adiposity [36, 37], and other adipose depots behave similarly. The longest timepoint examined was d28 p.i., at which immune cell numbers had not returned to baseline. Bona fide long-lived memory should be confirmed at extended intervals, ideally combined with fate-mapping given the marked tissue-specific differences in eosinophil longevity [24]. Obesity-induced WAT immune phenotypes have been reported to persist through weight loss [38], indicating that established dysfunction may not be fully reversible by metabolic correction alone, while even short-term HFD induces proliferation of memory adipose T cells and accumulation of a novel Th2-regulatory population [39]. Mechanistic resolution will benefit from interventional approaches such as weight-loss reversal, targeted cytokine neutralisation, and genetic manipulation of candidate niche components, and the human relevance of our findings remains to be tested in WAT samples from lean and obese individuals after natural helminth exposure or vaccination.

In summary, our findings establish white adipose tissue as a broadly competent type 2 immunological memory niche that accumulates both lymphoid and tissue-resident innate populations following pulmonary type 2 inflammation, and identify obesity as an important and selective disruptor of this depot. The apparent preservation of functional competence in cells that do establish provides a cellular framework for understanding the impaired vaccine responses and accelerated antibody waning associated with obesity across pathogens [20, 40], and positions WAT as a metabolically vulnerable component of the emerging paradigm of long-lived, epigenetically imprinted tissue type-2 immune memory [25].

## Methods

### Animals

Male C57BL/6J animals were purchased from Charles River (Sulzfeld, Germany) at the age of 6-8 weeks. *B6*.*SJL-Ptprc*^*a*^*Pepc*^*b*^*/BoyJ* (Ly5.1-B6) [41] mice were kindly provided by D. Voehringer (Universitätsklinikum Erlangen) and express the CD45.1-receptor variant on their immune cells. *C;129S4-Rag2*^*tm1*.*1Flv*^*Il2rg*^*tm1*.*1Flv*^*/J* (*Rag2*^*-/-*^ *Il2rg*^*-/-*^) [42] were kindly provided by U. Schleicher (Universitätsklinikum Erlangen). *Tg(TcraTcrb)425Cbn/J_CD45*.*1* (OT-II) [43] were kindly provided by C. Lehmann and D. Voehringer (Universitätsklinikum Erlangen). All genetically modified mice were backcrossed to C57BL/6 background for more than 12 generations. All animals were housed in a specific pathogen-free facility. Food and water were provided ad libitum and animal welfare was checked daily. All animal experiments were conducted in compliance with German animal protection law and approved by the Federal Government of Lower Franconia (RUF-55.2.2-2532-2-1177 and - 1813).

### Diet-induced obesity

Animals were fed a high-fat diet (HFD; metabolizable energy: 21.6 MJ/kg, 60% kcal fat, D12492; ssniff Spezialdiäten, Soest, Germany) or control diet (CD; metabolizable energy: 14.4 MJ/kg, 16% kcal fat; V1244-703; ssniff Spezialdiäten, Soest, Germany) for at least 12 weeks. Weight gain was monitored weekly.

### *Nippostrongylus brasiliensis* infection

Mice were infected by subcutaneous injection of 500 L3 larvae *N. brasiliensis* in 200 µl autoclaved H_2_O as previously described [44]. Mice were killed by anaesthetic overdose and analysed at indicated time points after infection. Lung worm burden was determined on day 2 p.i. by enumeration of emigrating larvae from minced lung [45].

### Papain treatment

Mice received 20 µg papain (Calbiochem, Cat# 5125) in 20 µl PBS by intranasal administration on three consecutive days. Mice were killed by anaesthetic overdose and analysed on indicated time points after treatment.

### Adoptive transfers

Splenic CD4^+^ T cells of OT-II_CD45.1 mice were isolated using magnetic activated cell sorting (MACS, MojoSort™ Mouse CD4 T cell Isolation Kit, Biolegend). Afterwards, 6 x 10^6^ CD4^+^ OT-II cells were intravenously transferred into lean or obese wild-type mice. One day later, recipient animals were infected with *N. brasiliensis* in combination with 100 µg ovalbumin (OVA). Therefore, 500 *N. brasiliensis* L3 larvae were incubated with 100 µg OVA for 30 minutes at 37°C under constant slow movement (New Brunswick Scientific, innova 2300 platform shaker; 50 rpm). Mice were killed by anaesthetic overdose and analysed 7 and 28 days post infection.

Ly5.1-B6 donor mice were kept on HFD or CD. After 12 weeks, mice were subcutaneously infected with *N. brasiliensis*. At 28 days post infection, gonadal WAT was extracted, digested and the stromal vascular fraction (SVF) from 2-3 mice was isolated by centrifugation and pooled for adoptive transfers. For CD4^+^ T cell transfers, T cells were magnetically isolated from the WAT (MojoSort™ Mouse CD4 T cell Isolation Kit, Biolegend), and 10^6^ cells were adoptively transferred intraperitoneally into *Rag2*^*-/-*^ *Il2rg*^*-/-*^ recipient mice. One day later, recipients were infected with *N. brasiliensis*.

### Sample preparation

Lung and WAT were minced into small pieces and digested using 1 mg/ml Collagenase D for 40 minutes at 37°C under constant shaking (New Brunswick Scientific, innova 2300 platform shaker; 120 rpm). The digested tissue was strained through a 100 µm cell strainer using a sterile syringe plunger and erythrocyte lysis was performed (ACK-lysis buffer: 150 mM NH_4_Cl, 10 mM KHCO_3_, 0.1 mM Na_2_EDTA in H_2_O, pH 7.2-7.4; 5 min, room temperature). Viable cells were counted in glasstic slides (KOVA) by trypan-blue (Sigma) exclusion of dead cells.

### Flow cytometry

2 × 10^6^ cells were transferred into V-bottom plates (Thermo Scientific) and stained with Zombie aqua fixable dye (BioLegend) in PBS for 20 minutes at 4°C. For surface staining, master mixes were prepared in FACS buffer (PBS, 2% FCS and NaN_3_) using the antibodies listed below, blocking of F_c_ receptors was performed by adding 10 µg/ml purified anti-CD16/32 antibodies (clone93; eBioscience). Staining was performed at 4°C for 40 min. Intracellular staining of transcription factors was performed using the FoxP3 staining buffer kit (eBioscience) following manufacturer’s instructions.

**Table 1:**
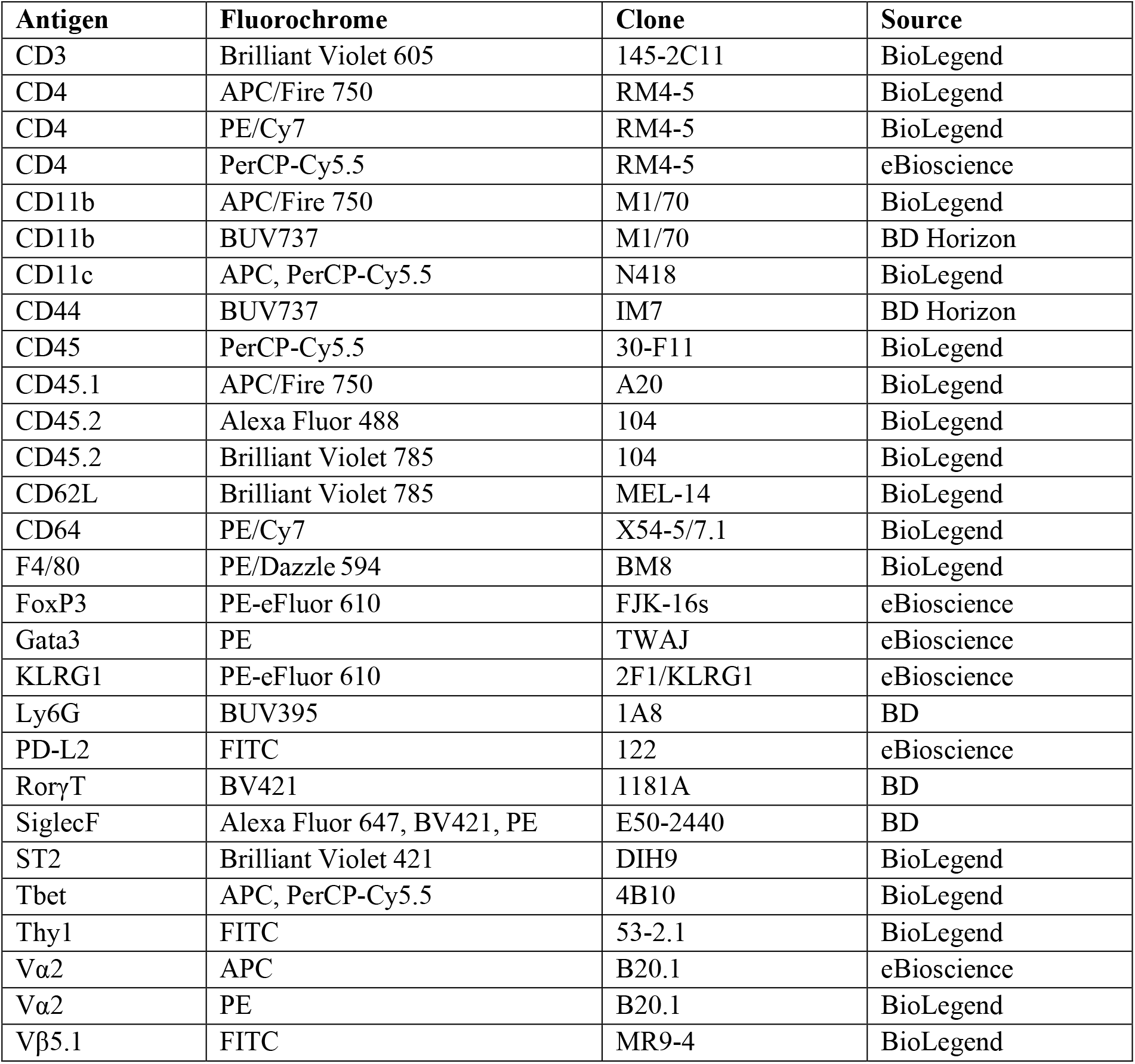
List of antibodies used in flow cytometry.

Eosinophils (CD45^+^CD11b^+^CD11c^-^SiglecF^+^), neutrophils (CD45^+^CD11b^+^Ly6G^+^), AAM (CD45^+^CD64^+^F4/80^+^PD-L2^+^), ILC2 (CD45^+^lin^-^Thy1^+^ST2^+^KLRG1^+^) were gated as described in Supplemental Fig. 1B, C. Naive T cells (CD4^+^CD44^-^CD62L^+^, effector/memory T cells (CD4^+^CD44^+^CD62L^-^), Th1 (CD4^+^Tbet^+^), Th2 (CD4^+^Gata3^+^) were gated as described in Supplemental Fig. 1D. OT-II cells were identified as CD4^+^CD3^+^CD45.2^-^CD45.1^+^Vα2^+^.

Samples were acquired on a BD LSR Fortessa flow cytometer. Fluorescence minus one (FMO) controls were used for gating. Data analysis was performed using FlowJo (version 10.10).

### Histology

The right lobe of the lung was used for histological analysis. Therefore, the lung lobe was kept in 4 % PFA directly after extraction at 4°C for at least 3 hours. Afterwards the PFA was removed and the lung was washed with PBS twice for another at least 2 hours. To stabilize the samples, lungs were then incubated in 10 % sucrose dissolved in PBS for 2 h, in 20 % sucrose for 2-4 h and in 30 % sucrose overnight. Every incubation step was performed at 4°C. Afterwards lungs were kept in a 1:1 dilution of 30 % sucrose and tissue freezing medium (Leica) for 2 h at room temperature (RT). To ensure the penetration of the freezing medium in the lung, the lungs were put into fresh freezing medium and a vacuum was admitted 3 times for 7 min with intermittent reoxygenation. The lungs were then put into cryomolds (Sakura) with fresh tissue freezing medium and after 1 h of incubation at RT, the organs were slowly frozen on dry ice. The samples were cryo-sectioned into 7 µm thick slices (CryoStar NX70 cryostat). Sections were collected on glass coverslips for haematoxylin and eosin (H&E) staining. Images were acquired on a Keyence BZ-X810.

### Post-acquisition and statistical analysis

Mice were randomly assigned to groups during the experiments. Flow cytometric data was analysed using FlowJo software (Treestar, v10.10). Statistical analyses were performed using GraphPad Prism (v9). Mean and SEM were calculated from repeated experiments. Outliers were identified by ROUT and normal distribution determined by Shapiro-Wilk normality test. Statistical significance was determined using one-way ANOVA and two-way ANOVA. Statistical significance was defined as *p*<0.05. In the figures, only relevant significances are depicted. All illustrations were created using BioRender.com. Figures were created using Affinity Designer (Serif Europe Ltd., v 3.0.1.3808).

## Data availability

The datasets generated during and analysed during the current study are available from the corresponding author upon reasonable request.

## Acknowledgments

L.S., A.D., I.H. and C.S. are supported by the Federal Ministry of Research, Technology and Space (BMFTR 01KI2109). L.S. and C.S. received support by the Deutsche Forschungsgemeinschaft (DFG, German Research Foundation; associated project within the Research Training Group “Immunomicrotope”, GRK 2740/447268119).

We thank C. Bogdan for his continuous support. Furthermore, we thank D. Voehringer for reagents, maintenance of the *N. brasiliensis* life cycle and mice, and U. Schleicher and C. Lehmann for reagents and mice. We thank M. Kirsch and the team of the Preclinical Experimental Animal Center (PETZ) for animal husbandry.

## Author Contributions

L.S. designed, performed and analysed experiments; drafted the manuscript. I.H. and A.D. contributed to specific experiments and edited the manuscript. C.S. conceptualised the study; designed and supervised the experiments; wrote the manuscript.

## Competing Interests

All authors declare no competing interests.

**Supplemental figure 1.**
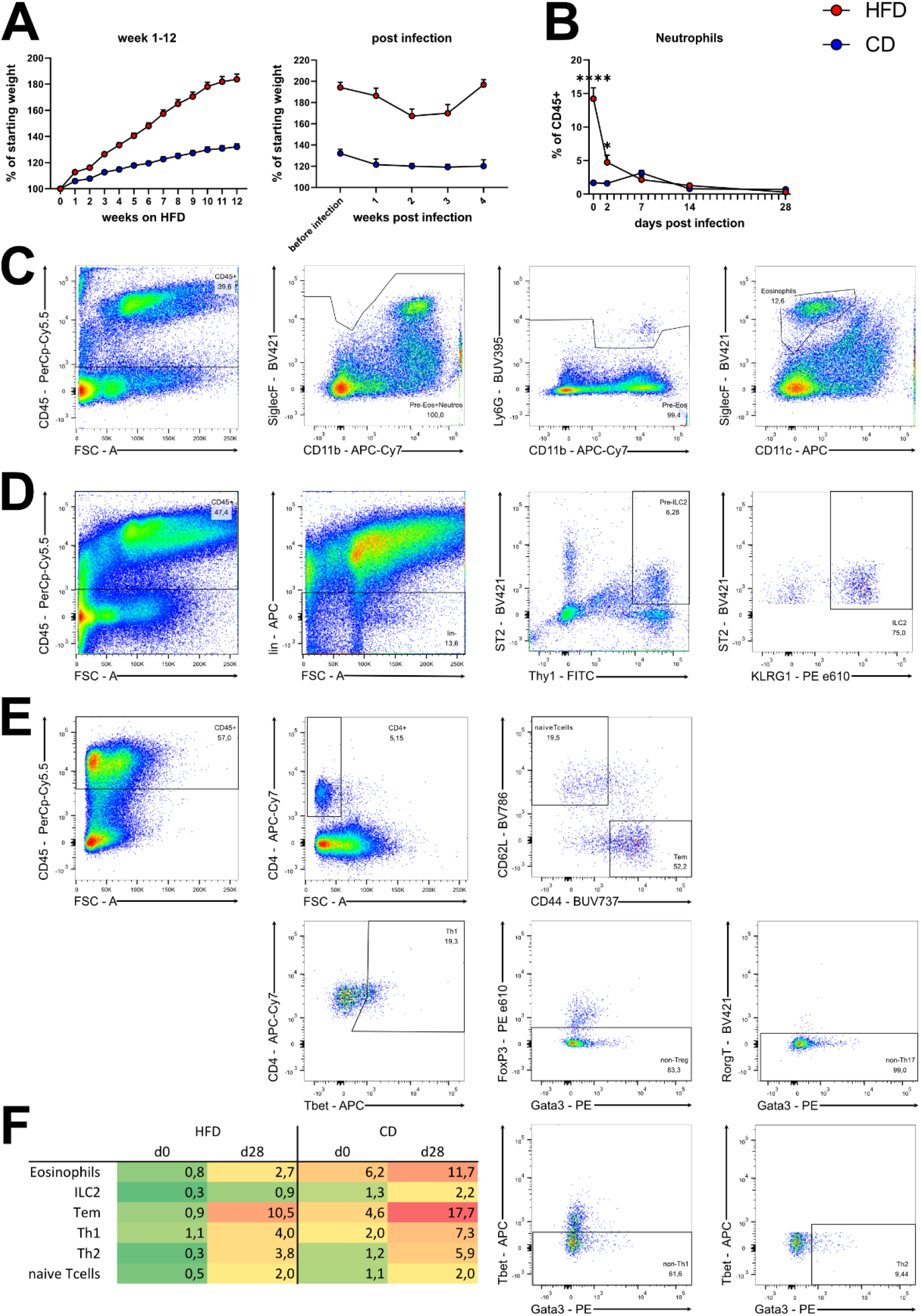
**A)** Body weight gain of mice kept on HFD (red) or CD (blue) for 12 weeks prior to experimental intervention. Graph shows mean + SEM of 20-34 mice per group from 3 independent experiments. **B)** Body weight change of mice kept on HFD (red) or CD (blue) and infected with *N. brasiliensis* during the first 4 weeks after infection. Graph shows mean + SEM of 16-27 animals per group from 3 independent experiments. **C**-**E)** Gating strategy to identify immune eosinophils **(C)**, ILC2 **(D)**, T cell subsets **(E)** in WAT.Pregated on living, single lymphocytes.**F)** Heatmap to visualize the frequency of highlighted cell populations in the WAT of naive (d0) and *N. brasiliensis* infected (d28 post infection) CD-fed and HFD-fed mice.

